# Unbiased stereological estimates of dopaminergic and GABAergic neurons in the A10, A9, and A8 subregions in the young male Macaque

**DOI:** 10.1101/2022.02.08.479596

**Authors:** Emily A. Kelly, Jancy Contreras, Annie Duan, Rochelle Vassell, Julie L. Fudge

## Abstract

The ventral midbrain is the primary source of dopamine- (DA) expressing neurons in most species. GABA-ergic and glutamatergic cell populations are intermixed among DA-expressing cells and purported to regulate both local and long-range dopamine neuron activity. Most work has been conducted in rodent models, however due to evolutionary expansion of the ventral midbrain in primates, the increased size and complexity of DA subpopulations warrants further investigation. Here, we quantified the number of DA neurons, and their GABA-ergic complement in classic DA cell groups A10 (midline ventral tegmental area nuclei [VTA] and parabrachial pigmented nucleus [PBP]), A9 (substantia nigra, pars compacta [SNc]) and A8 (retrorubral field [RRF]) in the macaque. Because the PBP is a disproportionately expanded feature of the A10 group, and has unique connectional features in monkeys, we analyzed A10 data by dividing it into ‘classic’ midline nuclei and the PBP. Unbiased stereology revealed total putative DA neuron counts to be 210,238 +/− 17,127 (A10 = 110,319 +/− 9,649, A9= 87,399 +/−7,751 and A8=12,520 +/− 827). Putative GABAergic neurons were fewer overall, and evenly dispersed across the DA subpopulations (GAD67= 71,215 +/− 5,663; A10=16,836 +/− 2,743; A9=24,855 +/− 3,144 and A8=12,633 +/− 3,557). Calculating the GAD67/TH ratio for each subregion revealed differential balances of these two cell types across the DA subregions. The A8 subregion had the highest complement of GAD67-positive neurons compared to TH-positive neurons (1:1), suggesting a potentially high capacity for GABAergic inhibition of DA output in this region.

**HIGHLIGHTS:**

- The A10 subregion expands laterally and caudally in nonhuman primates
- The A10, A9, and A8 comprise 52%, 42% and 6% of DA neurons, respectively
- GABAergic neurons are more evenly dispersed across subregions
- The A8 subpopulation has the highest ratio of GABA: DA neurons

## INTRODUCTION

An explosion of research in the last 20 years has significantly increased understanding of the function and diversity of the midbrain dopamine (DA) neuronal subregions, and their roles in behavior. These subregions--A10, A9, and A8--are commonly known as the ventral tegmental area (VTA), substantia nigra, pars compacta (SNc), and retrorubral field (RRF), respectively. It is now recognized that DA neurons’ role in diverse behaviors--from reward learning, to social motivation, to executive functions and motor output--results from a complex neuroanatomic and circuit organization through the midbrain, with differential gene expression profiles, electrophysiologic properties, and co-transmitter content (Watabe-Uchida et al., 2012; Beier et al., 2015; Lerner et al., 2015; Morales and Margolis, 2017; Zhang et al., 2017; Farassat et al., 2019; Poulin et al., 2020). For example, recent evidence, mainly from rodent models, indicates various DA neurons can co-contain glutamate (Sulzer et al., 1998; Root et al., 2016), and various neuropeptides (Seroogy et al., 1988; Bouarab et al., 2019; Pristera et al., 2019; Nagaeva et al., 2020).

Beyond DA neuron diversity, it has long been known that GABAergic neurons comprise the largest non-dopaminergic subpopulation in the ventral midbrain (Oertel et al., 1982; Nagai et al., 1983; Mugnaini and Oertel, 1985; Nair-Roberts et al., 2008; Margolis et al., 2012). GABAergic neurons are embedded among DA cells in each of the DA subregions. GABAergic neurons are also heterogeneous based on morphology and immunostaining for calcium binding proteins, peptides, and nitric oxide synthase (Parent et al., 1995; McRitchie et al., 1996; Paul et al., 2018; Nagaeva et al., 2020). Glutamatergic neurons also comprise a neuronal subpopulation in the ventral midbrain (Olson and Nestler, 2007; Nair-Roberts et al., 2008; Root et al., 2016), but are less abundant than GABAergic neurons and are largely confined to midline VTA (Nair-Roberts et al., 2008; Root et al., 2016). The non-dopaminergic cell types also have long-range projections, and it is not yet clear what proportion exert local control through axon collaterals, or simply function as interneurons (Van Bockstaele and Pickel, 1995; Carr and Sesack, 2000; Mailly et al., 2003; Omelchenko and Sesack, 2009; Dobi et al., 2010; Henny et al., 2012; Taylor et al., 2014).

DA neurons were classically divided into A10, A9, and A8 neuronal groups in rats and primates (Pearson et al., 1983; Bjorklund and Lindvall, 1984; Hokfelt et al., 1984), based on morphology and immunohistochemical staining to tyrosine hydroxylase (TH). Many of the chemical markers mentioned above have helped enormously to identify the neuronal groups across species. These studies show differences in size, shape and location of DA subregional boundaries, serving as cell-specific markers as the entire system expands from rodent to monkey and human (reviewed in, Bjorklund and Dunnett, 2007). The classic DA subgroup boundaries in monkey and human are drawn according to immunoreactivity for TH immunoreactivity, and for expression of the calcium binding protein D28k in TH neurons of specific subgroups (CaBP; a marker of the A10 and A8 DA subpopulations, but not the majority of A9 neurons) (Yamada et al., 1990; Lavoie and Parent, 1991; Gaspar et al., 1993; Haber et al., 1995; McRitchie et al., 1996). The A9 TH-IR neurons are specifically highly enriched in the g-coupled protein inward-rectifying potassium channel 2 (Girk2) (Schein et al., 1998; Chung et al., 2005; Margolis et al., 2012; Reyes et al., 2012; Fudge et al., 2017). Almost nothing is known of the relative distribution of GABAergic neurons within these subpopulations in human and nonhuman primates.

There are very few quantitative studies of the main cellular units in the DA subregions in nonhuman primates assessed by modern stereological techniques. To begin to understand to relative size and extent of the A10, A9, and A8 subregions, as well as the basic balance of DA and GABAergic cells within each, we applied stereological counting methods to the ventral midbrain regions in Macaques. Our goal was to develop a spatial map of the subregions in an animal model close to the human. We also sought to examine the complement of GABAergic neurons within and across the main DA subregions.

## EXPERIMENTAL PROCEDURES

### Animals

All experiments were carried out in accordance with National Institute of Health guidelines (NIH Publications No. 80-23). Experimental design and techniques were aimed at minimizing animal use and suffering and were reviewed by the University of Rochester Committee on Animal Research. To conserve animals, we used perfused tissue from 3 young male monkeys (*Macaque fascicularis*) that had received tracer injections in other brain regions as part of other studies (Table 1).

**Table 1.**
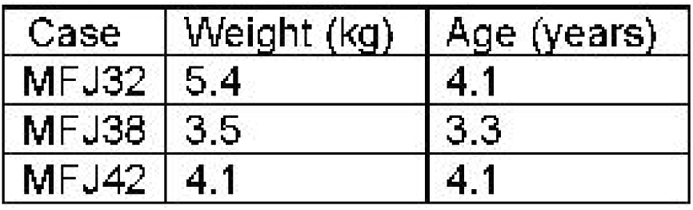
Experimental Cases.

| Case | Weight (kg) | Age (years) |
| --- | --- | --- |
| MFJ32 | 5.4 | 4.1 |
| MFJ38 | 3.5 | 3.3 |
| MFJ42 | 4.1 | 4.1 |

### Histology

All animals were sacrificed under deep anesthesia (pentobarbital) by intracardiac perfusion, first with 0.9% sterile saline and then 6L of 4% paraformaldehyde in a solution of sucrose and phosphate buffer. Brains were harvested and placed in 4% paraformaldehyde overnight, and then sunk in increasing gradients of sucrose solution (10%, 20%, and 30%). The entire brain was coronally sectioned on a freezing microtome at 40μm, and all sections were saved in serial wells (‘compartments’) in a cold cryoprotectant solution containing 30%sucrose and 30% ethylene glycol in phosphate buffer. They were stored at −20°C.

### Calbindin D28k (CaBP), glutamic acid decarboxylase 67 (GAD67), and tyrosine hydroxylase (TH) immunoreactivity (IR)

1:24 adjacent sections through the ventral midbrain were immunostained for TH (a synthetic enzyme, and marker, of DA neurons the ventral midbrain; Pearson et al., 1983), CaBP (a marker of the A10 and A8 DA subpopulations; Yamada et al., 1990; Lavoie and Parent, 1991; Gaspar et al., 1993; McRitchie et al., 1996; Haber and Fudge, 1997), and GAD67 (a synthetic enzyme for GABA, and marker of GABAergic neurons; Margolis et al., 2012). The A9 neuronal group is identified by its marked absence of CaBP-IR, which matches the distribution of relatively high Girk2-IR (not shown, Schein et al., 1998; Chung et al., 2005; Reyes et al., 2012; Fudge et al., 2017). Using 1:24 sections yielded 6 evenly spaced coronal sections through the ventral midbrain, with 960 μm between sections. In brief, we section the entire brain in the coronal plane, collecting each section in 24 consecutive wells. Therefore, each well has 1:24 sections with a unique rostrocaudal starting point, which is randomly selected for processing. After immunostaining the entire well, and mounting the tissue (see below), sections through the ventral midbrain region that contain the regions of interest are marked. Each case yielded 6 sections; therefore, the rostrocaudal extent of the ventral midbrain is approximately 5.75-6 mm.

Adjacent compartments through the ventral midbrain were processed using mouse anti-CaBP (1: 10,000, C9848, Sigma, St. Louis, MO), mouse anti-TH (1:10,000, MAB318, EMD Millipore Corp, Temecula, CA), and mouse anti-GAD67 (1:10,000, MAB5406, EMD Millipore Corp, Temecula, CA) (Table 1). These antibodies have been used in many studies across species (e.g. JCN Database 14.0), and their distribution was similar to that in recently published work on monkey and human (e.g. CaBP: Rice et al., 2016; eg. TH: Fudge et al., 2017; e.g. GAD67: Mabry et al., 2020; Selemon and Begovic, 2020). All cases (animals) for each individual stain were batch processed, under identical parameters. Selected sections were first thoroughly rinsed in 0.1M phosphate buffer solution containing 0.3% Triton-X (PB-TX) followed by an overnight incubation in the same solution The following day, tissue was treated with an endogenous peroxidase solution (10% methanol, 3% hydrogen peroxide [H_2_O_2_] in 0.1M PB pH 7.4) for 5 minutes and rinsed thoroughly in PB-TX. Sections were incubated in blocking solution containing 10% normal goat serum in PB-TX (NGS-PB-TX) for 30-min followed by a ∼96-hour shaker incubation in primary antisera and NGS-PB-TX at 4°C. Tissue was then rinsed in PB-TX, blocked with 10% NGS-PB-TX, and incubated in biotinylated goat anti-mouse secondary antibody (1:200, BA-9200, Vector Laboratories) for 40 minutes at room temperature. Following additional rinses, tissues were incubated for 1-hour in an avidin biotin complex solution (per kit instructions, ABC Peroxidase, PK-6100, Vector Laboratories) at room temperature. Tissues were then rinsed several times in 0.1M PB followed by 0.05M Tris Buffer (TB) and incubated in solution containing 0.05% 3,3’-diaminobenzidine (DAB) with 3% H_2_O_2_ in TB for 10 minutes. Sections were rinsed thoroughly in 0.1M PB and mounted out of a gelatin-based solution (0.05% 275 bloom gelatin, 20% ethanol in ddH_2_O) onto gelatin coated slides. Once dry, slides were dehydrated through xylene and cover-slipped using DPX mounting media (Electron microscopy systems [EMS], Cat. 13510).

Analysis

### Delineation of DA subpopulation boundaries (Fig. 1)

CaBP-IR of cell bodies was used to identify the general boundaries between the A10 neurons and A8 neurons, and the ventrally located A9 neurons (Lavoie and Parent, 1991). We also considered morphologic characteristics of TH-positive cells. The A10 neurons in the nonhuman primate extend the entire rostrocaudal extent of the midbrain, and include the following subnuclei: the rostral linear nucleus (RLi), the caudolinear nucleus (CLi), the intrafascicular nucleus, the ventral tegmental nucleus (VTA), the paranigral nucleus, and the parabrachial pigmented nucleus (PBP, Halliday and Tork, 1986; McRitchie et al., 1995). The PBP is an unusually large subnucleus of the A10 in the monkey and human (Halliday and Tork, 1986; Olszewski and Baxter, 2014), with large mediolaterally oriented neurons that sweep dorsally over the entire A9 region, and merge caudally with the A8 group. We therefore grouped all the VTA subnuclei medial to the PBP as the ‘midline VTA’ group and designated the PBP as a separate A10 subnucleus. The A9 neuronal group was delineated by the CaBP-poor region, which overlaps with Girk2-IR neurons (not shown, Fudge et al., 2017). The A9 DA subpopulation is also easily recognized with densely packed, large-sized TH-positive neurons (see Fig. 2) which form vertically oriented cell clusters with their dendrites reaching deeply into the pars reticulata. The A8 neurons are situated in the retrorubral field and are embedded in and around the fibers of the medial lemniscus. In coronal sections, at the level where the medial lemniscus is first detected, the medial A8 cell group merges with the caudal PBP.

**Figure 1.**
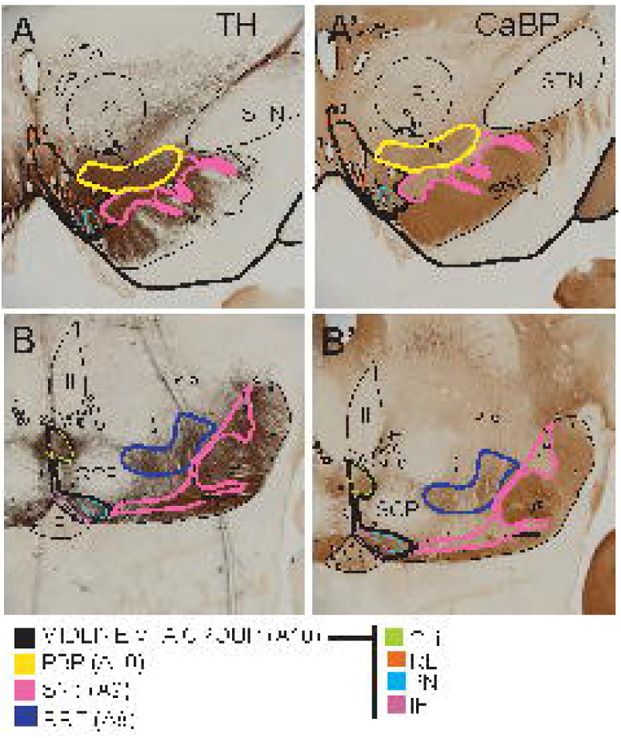
Macroscopic views of DA subpopulations in Macaque. A-B. TH-positive immunoreactivity in rostrocentral (A) and caudal (B) ventral midbrain. A’-B’. Adjacent sections immunostained for CaBP-protein. The A10 (black, midline VTA; yellow, PBP outlines) and A8 (blue outline) regions contain many CaBP-positive cell bodies, which are not appreciated at this magnification. The absence of CaBP-positive neurons is striking in the A9 subpopulation (pink outline). The nondopaminergic SNr has dense CaBP-positive neuropil. *Abbreviations: CLi, caudal linear nucleus; III, third nerve root; IF, intrafascicular nucleus; IP, interpeduncular nucleus; PBP, parabrachial pigmented nucleus; PN, paranigral nucleus; RN, red nucleus; RRF, retrorubral field; scp; decussation of the superior cerebellar peduncle; SNc, substantia nigra, pars compacta; SNr, substantia nigra, pars reticulata; STN, subthalamic nucleus; VTA, ventral tegmental area*.

**Figure 2.**
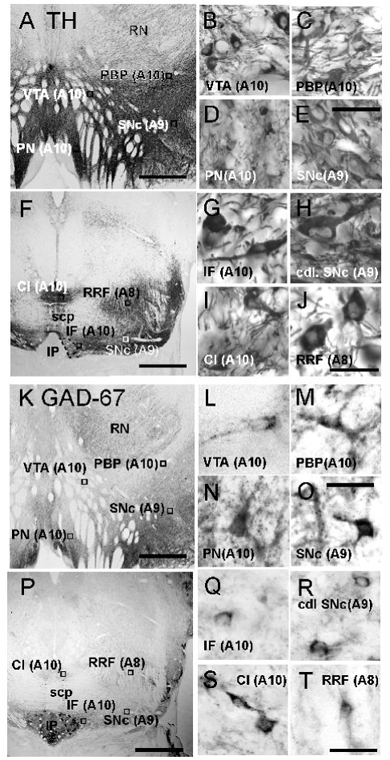
Representative examples of TH and GAD67 immunostaining. Photomicrographs through rostrocentral (A, K) and caudal (F, P) levels of the midbrain, depicting TH (A-J), and GAD67 (K-T) immunostaining in adjacent sections. Macroscopic images of TH immunostaining (A, F). Scale bar= 1 mm. Representative TH-positive neurons in selected regions are shown under higher power (B-E [rostrocentral], G-J [caudal]). Scale bar= 50μm. Macroscopic adjacent sections of GAD67 immunostaining (K, P). Scale bar= 1 mm. Representative higher power examples of GAD67-positive cells from boxed regions (L-O [rostrocentral],Q-T [caudal]). Scale bar= 50μm.

To define the boundaries of the subpopulations in each animal, sections were first photographed under low-power magnification (Plan Apo 2X/ 0.08 objective, Olympus BX51 brightfield microscope equipped with an Insight 4p Mosaic camera) and saved as a TIFF image using the SPOT 5.6 software (v. 5.6.3, Spot Imaging, Diagnostic Instruments, Inc.). Boundaries for the subnuclei and cell groups were mapped onto the TIFF image at the microscope under higher power (UPlan 10X/ 0.30 objective), with reference to cellular morphology and location.

### Unbiased stereology of TH-positive and GAD67-positive neurons

In conjunction with subregion maps (Fig. 1), we used the optical fractionator method (West and Gundersen, 1990) to count neurons in the A10 ‘midline VTA’ and the PBP, the A9 group, and the A8 group. Slides were examined using an Olympus Provis AX70 microscope equipped with a Prior motorized stage (Proscan III) and mounted with an Olympus DC1394 (IIDC) camera under brightfield conditions. Sampling was implemented using the Microbrightfield StereoInvestigator software package (v. 2021.1.3, MBF Bioscience, Williston, VT) at 40X magnification (UPlanFI 40x/0.75 air objective). For GAD67-IR sections, we focused only on the A10, A9, and A8 subpopulations; therefore, the rostromedial tegmental nucleus (RMTg), pars reticulata, and the interpeduncular nucleus (IP), were excluded from the analysis. Neurons in the animal’s left hemisphere were counted. Subpopulation boundaries were registered onto TH-IR sections and adjacent GAD67-IR sections in the same animal, aligning fiducial markers such as blood vessels in the adjacent sections. Prior to the beginning of the study, average section thickness was measured on several sections from different brains, resulting in an approximate 25-30 μm final section thickness. Due to differences in cell size and packing densities across the subpopulations, sampling parameters for TH- and GAD67-IR neurons in each subregion were determined in advance in pilot studies. In all cases, we chose parameters where the Gundersen coefficient of error, m=1 (CE), was <0.1. For all sections, we selected a 15 μm Z plane to avoid any top or bottom cell distortions, with a 5 μm guard zone. Based on preliminary studies for each region, final sampling parameters for TH-IR sections were: PBP, A8, SNc: a 100 × 100 counting frame, and grid size of 212 × 212 was selected; midline VTA: 100 × 100 counting frame, and grid size 300 × 300. For GAD67-IR sections, sampling parameters were: PBP, A8, SNc: 100 × 100 counting frame with 115 × 115 grid size; midline VTA nuclei: 115 × 115 frame and 230 × 230 grid size. Section height was measured at each sampling site, and the estimated population was calculated using number weighted section thickness. The top of each cell was chosen as the ‘leading edge’ for counting. The total neuron number for each cell type in each region is estimated by the equation N=ΣQ × 1/ssf × 1/asf × 1/hsf, where ΣQ is the actual cell number counted in the sample, ssf= section sampling fraction (sampling interval, 1:24), asf= area sampling fraction (grid size/counting frame area), and hsf= height sampling fraction (average mounted section thickness/dissector height).

### Statistics

Unbiased, blinded counts of TH- and GAD67-positive neurons in each region of interest were summed for each case and compared across animals using a Mann Whitney test (pooled estimates) and a two-way ANOVA with Tukey multiple comparisons test (corrected) to investigate population estimates across ventral midbrain regions. Ratios of TH/GAD67-positive neurons per region was also calculated, using sums of cells in each region across all three animals. The relative proportion of TH and GAD-67-IR neurons in each section through the anterior-posterior plane were also calculated for each region. Estimated cell counts per level in each of the cases were pooled, and presented as a log10 scale for presentation/comparison purposes. A two-way ANOVA with Tukey’s multiple comparison tests were applied to assess rostrocaudal differences across cell-type at each rostrocaudal level (S1-S6). p<0.05=*, p<0.01=** (#), p<0.001=*** (∞), p<0.0001<=**** (+). Error bars are presented as SEM.

## RESULTS

### The A10, A9, and A8 subpopulations

The boundaries of A10 (VTA, PBP), A9, and A8 neuronal groups at two rostrocaudal levels of the midbrain are shown in Fig. 1. The midline VTA subnuclei boundaries are shown in black, the PBP in yellow, the SNc in pink, and the RRF in blue. The TH-IR neurons across the ventral midbrain vary in size, orientation and morphology as has been reported previously (Yelnik et al., 1987; Fig. 2A-J). The TH-containing neurons in midline VTA nuclei were generally smaller than those in the PBP, SNc, and RRF (Fig 2D compared to B, C and E-F). There was morphologic heterogeneity with paranigral TH-positive neurons being very small and round (15-20 μm), while neurons in other regions (i.e the VTA, Rli and Cl subnuclei) were slightly larger and often bipolar. TH-positive neurons in the PBP, SNc, and RRF were often very large, with elongated soma measuring approximately 40-50 μm in diameter, flanked by thick, tapering proximal dendrites that were oriented in the same plane.

GAD67-positive neurons were present and interspersed across the A10, A9, and A8 regions. These cells were much more diffusely distributed than TH-positive neurons, and had small round, or multipolar soma. Across subregions, GAD-67-positive neurons ranged from approximately 15-30 μm in diameter and had varying intensities of GAD67 immunoreactivity in all regions.

### Dopaminergic neuron counts

The total number of DA neurons per hemisphere, indicated by TH-labeling, is shown in Table 2. Per hemisphere, the average number of TH-positive cells for all subpopulations (n=3, 3 subpopulations) was 210,238 +/− 17,127, consistent with prior studies in monkey (Blesa et al., 2012; Selemon and Begovic, 2020). The total A10 neurons comprised 52% of the total DA population (110,319 +/− 9,649), with the midline VTA nuclei containing 29% and the PBP containing 23% of total TH-IR cells. The A9 and A8 accounted for 42% (87,399 +/− 7,751) and 6% (12,520 +/− 827) of total TH-IR cells, respectively.

**Table 2.** Estimates of TH- and GAD67-positive neurons for each subregion in all cases. The midline VTA and PBP comprise the A10 subregions, the SNc is the A9 subregion, and the RRF is the A8 subregion.

| REGION | CASE | TH | GAD-67 |
| --- | --- | --- | --- |
| MIDLINE<br>VTA(A10) | J32 | 58,588 | 21,832 |
|  | J38 | 53,262 | 25,678 |
|  | J42 | 71,809 | 30,732 |
|  | midline VTA-only Total | 184,657 | 88,043 |
| PBP (A10) | J32 | 34,840 | 13,344 |
|  | J38 | 55,860 | 10,575 |
|  | J42 | 55,700 | 9,218 |
|  | PBP-only Total | 146,299 | 33,136 |
| Total |  | 330,957 | 121,182 |
| Mean ± SEM | | 110,319 ± 4,669 | 33,727 ± 1,925 |
| SNc (A9) | J32 | 77,744 | 18,961 |
|  | J38 | 81,725 | 25,802 |
|  | J42 | 102,729 | 28,701 |
|  | Total | 262,198 | 74,465 |
| Mean ± SEM | | 87,399 ± 7,751 | 24,855 ± 3,148 |
| RRF (A8) | J32 | 15,732 | 8,652 |
|  | J38 | 10,940 | 18,731 |
|  | J42 | 12,888 | 9,018 |
|  | Total | 39,560 | 37,400 |
| Mean ± SEM | | 13,187 ± 827 | 12,467 ± 1,337 |
| Total Mean ± SEM | | 210,298 ± 17,127 | 71,215 ± 6,683 |

### GABAergic neuron counts

GAD67-positive neurons across the A10, A9, and A8 regions were summed and averaged across cases (Table 2, n=3). The substantia nigra pars reticulata and RMTg, which are GABAergic, and outside the DA subregions, are excluded from these analyses. Overall, there were 71,215 +/− 5,663 GAD67+ neurons among the A10, A9, and A8 subpopulations per hemisphere. These were found in all the DA subregions, consistent with rodent studies (Nair-Roberts et al., 2008). The mean GAD67-labeled cells in the A10 (uniliateral) were 33,727 +/− 1,923 (47% of the total GAD67 population [32% for the midline VTA nuclei and 15% for the PBP]). The A9 GAD67-IR neurons totaled 24,855 +/− 3,144 (35% of the total), and the A8 GAD67 neurons were 12,633 +/− 3,557 (18% of the total).

Estimated counts of each type of neuron were also assessed across individual cases (Table 3) to ascertain within-animal variability. For all animals, there was a significant difference in between cell-type comparisons only (TH-positive vs GAD67-positive, two-way ANOVA with Tukey’s multiple comparison, F(1,2)= 62.89, p=0.0155), and no significant differences in the number of each cell type across cases [F(2,2)=1.118, p=0.4722]). Thus, there was not significant individual variability across the three male animals of approximately the same age and pubertal status.

**Table 3.**
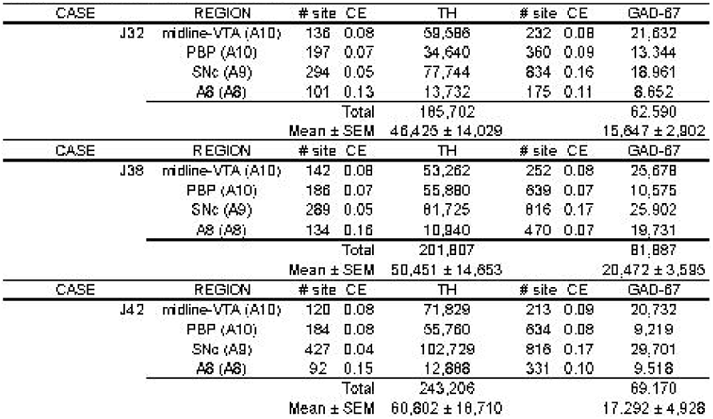
Estimates of TH-positive and GAD67-positive neurons across experimental cases.

### Proportions of TH- and GAD67-positive neurons by subregion

There were significantly more TH-positive cells compared to GAD67-positive cells overall (Table 2, Fig. 3A, Mann Whitney test, p=0.05). Overall, there were approximately 3 times more TH-positive than GAD67-positive neurons in the ventral midbrain as a whole. By subregion, significant differences in TH-IR cell numbers were found between the midline VTA group (triangles) and SNc and RRF, but not between the midline VTA and PBP (both A10 groups) (two-way ANOVA with Tukey’s multiple comparisons test, F(3,16)=16.11, p<0.0001; VTA vs SNc, p=0.0178, VTA vs RRF, p<0.0001, VTA vs PBP, p=0.5205). Similarly, the PBP (gray circles) had significant differences in TH-positive cell numbers compared with the SNc (open circles, Tukey’s multiple comparisons test, p=0.0004) and RRF (black squares, Tukey’s multiple comparisons test, p=0.0008). The SNc had significantly greater TH-IR cell counts than all other regions (Tukey’s multiple comparisons test; midline VTA [p=0.0178], PBP [p=0.0004], RRF [p<0.0001]), while the RRF had significantly fewer TH-positive neurons than all other regions (Tukey’s multiple comparisons test; midline VTA [p<0.0001], PBP [p=0.0008], and SNc [p<0.0001]).

**Figure 3.**
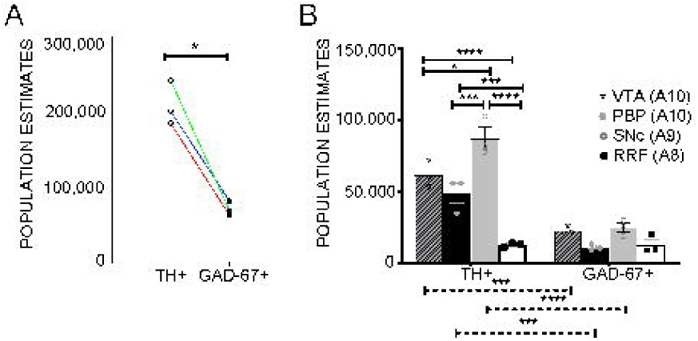
Estimated numbers of TH-positive and GAD67-positive cells in primate ventral midbrain. (A) Optical fractionator estimates of *total* TH-positive (open circles) and GAD67-positive (closed circles) cell counts depicting a significant 3-fold more TH-positive cells throughout the ventral midbrain. Counts were pooled across n=3 cases. Line colors denote each case. Mann-Whitney test, p=0.05. (B) Stereological estimates of cell types in A10 (VTA + PBP), A9 (SNc) and A8 (RRF) subregions. A significant proportion of cells originated from the midline VTA (triangle, hatched bar), PBP (gray circles, black bar) and SNc (open circles, gray bar) compared to the RRF (black squares, white bar). Solid significance bars correspond to “within cell type” comparisons. Two-way ANOVA with Tukey’s multiple comparison tests, F(3,16)= 16.11, p<0.001. Dashed significance bars correspond to “across cell type” comparisons denoting significantly more TH-positive cells compared to GAD67-positive cells in all groups except in the RRF.

When compared across DA subregions, there were no significant differences in GAD67-positive neurons in the midline VTA, the PBP, SNc, or RRF. Comparing TH-IR and GAD-67-IR neuron numbers across subpopulations, there were significantly more TH-IR neurons compared to GAD67-positive cells for every subpopulation except the RRF (A8) (Fig. 3B, two-way ANOVA with Tukey’s multiple comparisons test, solid black pairwise comparison lines, F(3,16)=16.11, p<0.001). GAD67-positive cell counts did not differ significantly across subregions.

We then calculated the ratio of GAD67 to TH-labeled cells for each subregion, to compute the relative contribution of each cell type within the classic DA subpopulations. This revealed that the A8 (RRF) subpopulation had the highest complement of GAD67-positive neurons, slightly exceeding the proportion of TH-positive cells (Table 4). In contrast, all other subpopulations had a significantly lower relative proportion of GAD-67-positive neurons compared to TH-IR cells. The proportion of GAD-67-IR neurons, compared to TH-IR neurons, in the midline VTA A10 was 38%, and 25% and 28% in the PBP and SNc, respectively.

**Table 4.** TH-positive/GAD67-positive and GAD67-positive/TH-positive ratios.

| REGION |  | TH/GAD RATIO | GAD/TH RATIO |
| --- | --- | --- | --- |
| MIDLINE VTA (A10) | J32 | 2.75 | 0.36 |
|  | J38 | 2.07 | 0.48 |
|  | J42 | 3.46 | 0.29 |
| | | $2.76 \pm 0.40$ | $0.38 \pm 0.06$ |
| PBP (A10) | J32 | 2.60 | 0.39 |
|  | J38 | 5.28 | 0.19 |
|  | J42 | 6.05 | 0.17 |
| | | $4.64 \pm 1.05$ | $0.25 \pm 0.07$ |
| SNc (A9) | J32 | 4.10 | 0.24 |
|  | J38 | 3.16 | 0.32 |
|  | J42 | 3.46 | 0.29 |
| | | $3.57 \pm 0.28$ | $0.28 \pm 0.02$ |
| RRF (A8) | J32 | 1.59 | 0.63 |
|  | J38 | 0.55 | 1.80 |
|  | J42 | 1.35 | 0.74 |
| | | $1.17 \pm 0.313$ | $1.06 \pm 0.374$ |

### Rostrocaudal shifts for TH and GAD67

We also evaluated the relationship between TH-positive and GAD67-positive neurons along the rostro-caudal axis (Fig. 4A-F). Levels are arranged from the rostral-most section (S1) of the ventral midbrain at the level of the subthalamic nucleus (STN), through the last section of the RRF (S6), at the level of the interpeduncular nucleus (IP). The RRF occupies only the latter third of the ventral midbrain (levels S5 and S6; Fig. 4E-F). Level designations are relative, and not aligned to a specific stereotaxic coordinate since sections were selected at random starting points for purposes of stereology. TH-IR neuron counts generally exceeded GAD67-IR neuron counts at most levels for the midline VTA, PBP and SNc (Fig. 4 G-I; see Table 5 for estimated count data), but converged in the caudal midbrain where TH-IR and GAD-67-IR cell counts were most similar to one another (Fig. 4J; see Table 5 for estimated count data). To investigate potential differences at each rostrocaudal level, we performed individual two-way ANOVAs to assess the effect of dopaminergic subregeion (VTA, PBP, SNc and RRF) and cell type (TH vs GAD67) on estimated neuronal counts (Table 6). We found a significant difference between cell types at several analyzed levels (specifically S2-S4, see Table 6 for statistical results). We also noted a significance based on the particular subregion at the S1 and S3 levels. To determine which rostocaudal level might be influencing the overall neuronal estimates the most, we further performed Tukey’s mulitple comparisons tests amongst the estimated number of dopaminergic (TH+) and GABAergic (GAD67+) neurons within each subregion and at each rostrocaudal level. Here we noted significantly more TH-positive (black circle) to GAD67-positive (gray circle) at S2 (in VTA, p=0.0059; in SNc, p=0.0353), S3 (in VTA, p=0.0118; in PBP, p=0.0002; and SNc, p=<0.0001) and S4 (in SNc, p=0.0052). Estimated count differences between TH-positive and GAD67-positive neurons at the most rostral and caudal levels were not significantly different.

**Figure 4.**
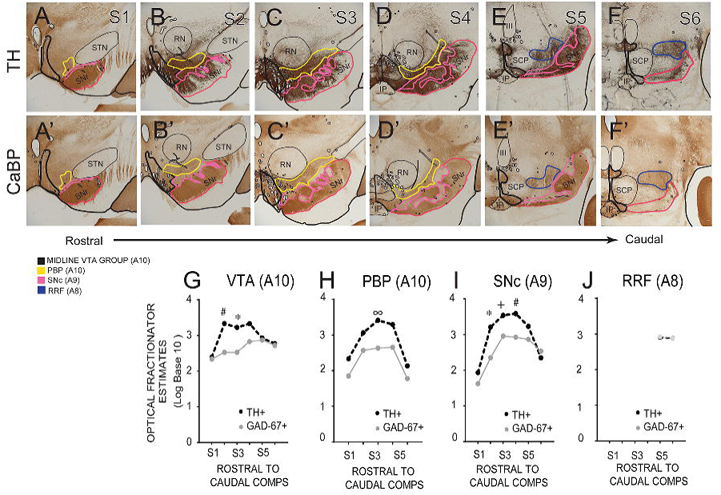
Rostrocaudal comparisons. Photomicrographs at evenly spaced intervals through the ventral midbrain, with DA subregions drawn in (A-F; case MF32) assisted by CaBP-IR in adjacent sections (A’-F’) along the rostrocaudal axis. Midline VTA groups (black outline), PBP (yellow outline), SNc (pink outline) and RRF (blue outline) are denoted at each level. G-J. Rostral (S1) to caudal (S6) TH-positive and GAD67-positive estimated cell counts in midline VTA (G), PBP (H), SNc (I) and RRF (J). Optical fractionator estimates are expressed as a Log Base 10 scale for presentation purposes (see Table 5 for estimated count data). We noted significant differences in estimated TH+ vs GAD67+ neurons at several levels. Two-way ANOVA with Tukey’s multiple comparisons tests. *= p<0.05, #= p<0.01, ∞= p<0.0005, += p<0.0001.

**Table 5.**
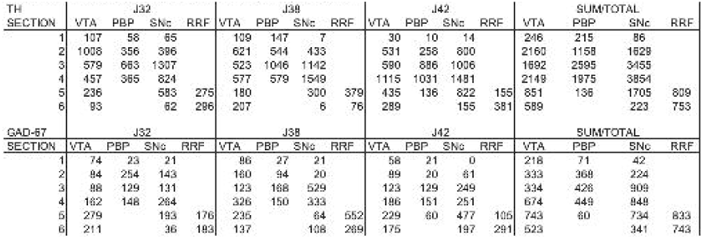
Rostrocaudal comparisons of TH-positive and GAD67-positive estimated cell counts across experimental cases.

**Table 6.** Two-way ANOVA results of rostral to caudal estimated neuron counts.

| **Level** | **S1** | **S2** | **S3** |
| --- | --- | --- | --- |
| **Cell Type** | F(1,12)=2.608 (p=0.1323) | F(1,12)=34.73 (p<0.0001)*** | F (1, 12) =118.6 (p<0.0001)*** |
| **Subregion** | F(2,12)= 3.970 (p=0.0475)* | F(2,12)=1.558 (p=0.2503) | F (2, 12) =13.26 (P=0.0009)** |

| **Level** | **S4** | **S5** | **S6** |
| --- | --- | --- | --- |
| **Cell Type** | F (1, 12) =29.44 (p=0002) ** | F (1, 12) = 1.359 (p=0.2664) | F(1,8)=1.125 (p=0.3197) |
| **Subregion** | F (2, 12) =3.623 (p=0.0588) | F (3,12) = 1.586 (p=0.2443) | F(1,8)=3.491 (p=0.0986) |

## DISCUSSION

An explosion of studies in recent years documents profound heterogeneity of DA cells across the expanse of the ventral midbrain, which includes circuit-specific differences in neurochemical and electrophysiologic properties of specific DA neurons. However, basic data on the anatomy of the DA subregions is lacking in the Macaque, a model which is the best anatomic match to the human. Diseases associated with DA neuron dysregulation or pathology are well-described in humans, ranging from mood disorders, social anxiety disorders, drug addictions and schizophrenia to neurodegenerative conditions such as Parkinson disease. As such, precision in mapping circuit specific paths and functions in a higher animal model is needed. We recently showed subregion-specific inputs/outputs in amygdala-DA-striatal paths in the Macaque, which involved the PBP and A8 neurons (Fudge et al., 2017). Meanwhile, behavioral roles for projection-specific DA cells inside and outside the midline VTA have now been identified in monkeys, using electrophysiologic techniques. These studies show that the anatomic position of the DA neurons predict specific circuits and functions (Kim et al., 2014, 2015; Stauffer et al., 2016). In this study, we take a first step in quantifying the distribution of DA neurons and their GABAergic counterparts throughout the A10, A9 and A8 subregions of the ventral midbrain in the macaque. We found significant subpopulation differences in the relative balance of putative DA and GABAergic neurons, which will inform future anatomic and physiologic studies.

### Comparison with previous studies in nonhuman primates

Our stereological results for total TH-IR neuron counts are generally consistent with several other nonhuman primate studies using similar stereological techniques (Emborg et al., 1998; Francois et al., 1999; Blesa et al., 2012; Selemon and Begovic, 2020). For comparison, in humans, TH-positive neuron counts are about twice the number found in the nonhuman primate, although this literature is widely divergent depending on methods (reviewed in Stark and Pakkenberg, 2004; Bjorklund and Dunnett, 2007; Fu et al., 2016). Previous studies in nonhuman primates did not divide the TH-neurons into subregions prior to sampling and used uniform sampling parameters throughout the entire structure. Most did not take the RRF (A8) group into account (although see Francois et al., 1999). Our study represents an attempt to apply standard boundaries for each subpopulation based on calbindin-D28K staining in DA neurons, which is a reliable marker across species (McRitchie et al., 1996; Fu et al., 2016). We also updated subdivisional schemes based on newer appreciation of the DA subregions in the nonhuman primate. Important to note is the appreciation of the parabrachial pigmented nucleus (PBP) as a lateral extension of the classic A10 VTA (Halliday and Tork, 1986; Olszewski and Baxter, 2014). Prior to widespread use of subregional immunocytochemical markers, this region was considered to comprise the ‘dorsal A9’ group by many groups, including ours (Francois et al., 1999; Cho and Fudge, 2010). Older studies therefore likely over-represent A9 counts by including the PBP in the dorsal A9.

There are no studies, to our knowledge, quantifying GAD67-positive counts throughout the full extent of the classic DA subpopulations in Macaque. Our finding that there are about 3 times more DA neurons than GABAergic neurons in the DA subpopulations is consistent with studies in rats (Swanson, 1982; Nair-Roberts et al., 2008). A study using in situ hybridization for GAD65/67 in rat compared percentages of putative GABAergic neurons to total neurons (labeled by NeuN), and found that GAD65/67 mRNA positive neurons formed approximately 23% and 13%, respectively, of total neurons in the VTA and SNc, respectively (compared with TH-immunoreactive cells comprising 55% and 88% of total neurons, in the same regions) (Margolis et al., 2012).

In mice, GABAergic neuron heterogeneity can be extensively studied in transgenic animals, as well as in different strains (reviewed in Bouarab et al., 2019), but to our knowledge, GAD67-labeled cell numbers have not directly been compared to TH-positive neuron counts in this species using modern cell counting techniques. An important endeavor for all animal models is to attempt to define specific GABAergic groups as interneurons versus long-range projection neurons using co-markers (Paul et al., 2018). In mice, optogenetic and chemogenetic techniques can enable silencing of long-range GABAergic projections, and thus dissociate interneuron activity and function (Tan et al., 2012; van Zessen et al., 2012). It will be important to determine the proportion of GABAergic neurons that actually participate in long-range projections in monkeys, especially with respect to anatomic location in the midbrain.

### Species variations highlight the need to expand knowledge in monkeys

The complexities of the ventral midbrain subregions differ across species (Bjorklund and Dunnett, 2007), in both the nature of efferent/afferent connectivity as well as cellular composition. The ventral midbrain of rats has about 20,000-30,000 TH-positive neurons per hemisphere (German and Manaye, 1993); mice have approximately 10,000-15,000 depending on strain (Nelson et al., 1996; Zaborszky and Vadasz, 2001; Brichta and Greengard, 2014). The ventral tegmental area (VTA) is a primary source of dopamine to the cortex, ventral striatum, and amygdala in the rodent. Newer studies in rodents make clear that DA neurons in the medial versus ‘lateral’ VTA have differential projections and responses to ‘emotional cues’ (i.e.Lammel et al., 2011; Chaudhury et al., 2013; Beier et al., 2015; Lerner et al., 2015; Morales and Margolis, 2017).

In our own work examining connections of the midbrain DA system, we have found that the midline VTA (A10) and lateral A10 (PBP) have differential projections to the striatum and amygdala. As a group, the midbrain DA neurons project to the striatum in progressive increments along their mediolateral and dorsoventral axis, to target the striatum in a ventromedial to dorsolateral gradient (Haber et al., 2000). With respect to the amygdala, we have found that the midline VTA is not the sole, or main input projection; rather the majority of DA-amygdala projections originate in the A10 neurons lateral to the midline VTA -i.e. the PBP (previously termed the A9dt)-- and the A8 neurons (Cho and Fudge, 2010). Our recent work indicates that, in the opposite direction, the amygdala-DA pathway also largely terminates outside the midline VTA, with labeled terminals most concentrated over the A10 PBP, A9 dorsal SNc, and the A8 RRF (Fudge et al., 2017). In other work, discrepancies regarding dopaminergic projection patterns to the cerebral cortex have been shown between rodent and higher order primates (Lewis et al., 1987; Berger et al., 1991). In primates, large regions of the cortex (including the motor, supplementary motor, parietal, and association cortices) receive DA innervation, whereas in rodents, DA terminals are largely confined to the anterior cingulate and to a lesser extent. Indeed, older tract tracing studies which apply large retrograde injections into various cortical regions in nonhuman primates indicate that much of the cortical input from the ventral midbrain originates in a broad continuum of midbrain neurons stretching from the midline VTA, across the ‘dorsal A9’ (now recognized as the PBP) to the A8 RRF (Gaspar et al., 1989; Williams and Goldman-Rakic, 1998).

### Functional Significance

It is now well-established that the DA system can no longer be considered as simply a ‘two transmitter system’ composed of either DA or GABAergic neurons. Cells throughout the DA subpopulations are an admixture of DA-, glutamate-, and GABA-containing neurons (Dobi et al., 2010; Margolis et al., 2012; Yamaguchi et al., 2015; Root et al., 2016; Morales and Margolis, 2017), a proportion of which are ‘multiplexed’ (expressing multiple neurotransmitter profiles), and participate in a variety of motivated behaviors (Lammel et al., 2012; Tan et al., 2012; Root et al., 2018; Farassat et al., 2019). A large amount of recent work has focused on the complex role of glutamatergic signaling in the ventral midbrain mainly in the rodent (Dobi et al., 2010; Gorelova et al., 2012; Hnasko et al., 2012; Morales and Margolis, 2017; Root et al., 2018), but more recently, in non-human primates as well (Root et al., 2016).The anatomic and functional role of GABA, amongst and within these cell groups, is less understood (see review by, Bouarab et al., 2019). VTA GABAergic neurons drive conditioned place aversion (Tan et al., 2012) and disruption of reward consumption (van Zessen et al., 2012; Wakabayashi et al., 2019; Lowes et al., 2021), and their activation is associated with exposure to acute stressors (Cohen et al., 2012; Tan et al., 2012; Zhou et al., 2019). Conversely, inhibiting GABAergic neurons in the ventral midbrain increases DA firing and is associated with opiate reward-seeking (Margolis et al., 2014; Galaj et al., 2020). While GABAergic activation can inhibit DA firing (at least in the VTA), the question of whether this occurs via direct interneuron-like activity, or a less direct route is still not clear. Given these basic questions, determining the complement of DA and GABAergic neurons in anatomic subpopulations of primate ventral midbrain is an important starting point.

While some of our general results were similar to existing rodent data, this was not the case when we examined the relationship of GABA:DA neuron in the RRF (A8). In contrast to other regions where the proportion of GABAergic neurons were significantly less than DA cells, an equal proportion of each neuron type occurred in the A8 RRF. Functionally, A8 falls within the presumptive “salience-detecting” neurons (Matsumoto and Hikosaka, 2009; Matsumoto and Takada, 2013). In the nonhuman primate, these neurons project to the ‘limbic associative striatum’, (reviewed in, Francois et al., 1999; Haber, 2014; Fudge et al., 2017), the amygdala (Cho and Fudge, 2010), the thalamus (Sanchez-Gonzalez et al., 2005), and broadly throughout the cortex (Gaspar et al., 1989; Williams and Goldman-Rakic, 1998). The 1:1 relationship between TH and GAD67 (presumptive elevated inhibitory tone) within this region may suggest differential modulation of dopaminergic signaling, or even the presence of long-range GABA projections, to these downstream targets.

## ABBREVIATIONS

CaBP: calcium binding protein D28k
CLi: caudal linear nucleus
DA: dopamine
III: third nerve root
GABA: Gamma-Aminobutyric Acid
GAD67: glutamic acid decarboxylase 67
Glu: Glutamate
IF: intrafascicular nucleus
IP: interpeduncular nucleus
PBP: parabrachial pigmented nucleus
PN: paranigral nucleus
RN: red nucleus
RRF: retrorubral field
RMTg: rostromedial tegmental nucleus
scp: decussation of the superior cerebellar peduncle
SN: substantia nigra
SNc: substantia nigra, pars compacta
SNr: substantia nigra, pars reticulata
STN: subthalamic nucleus
TH: tyrosine hydroxylase
VTA: ventral tegmental area.

## Acknowledgements

This work was supported by the National Institutes of Mental Health (grant number R01MH115016).

## REFERENCES CITED

Beier KT et al. (2015) Circuit architecture of vta dopamine neurons revealed by systematic input-output mapping. Cell 162:622–634.

Berger B, Gaspar P, Verney C (1991) Dopaminergic innervation of the cerebral cortex: Unexpected differences between rodents and primates [published erratum appears in trends neurosci 1991 mar;14(3):119]. Trends in Neurosciences 14:21–27.

Bjorklund A, Lindvall O (1984) Dopamine-containing systems in the cns. In: Handbook of chemical neuroanatomy, vol. Ii: Classical transmitters in the cns, part i (Bjorklund, Hokfelt, eds), pp 55–122. Amsterdam: Elsevier.

Bjorklund A, Dunnett SB (2007) Dopamine neuron systems in the brain: An update. Trends Neurosci 30:194–202.

Blesa J et al. (2012) The nigrostriatal system in the presymptomatic and symptomatic stages in the mptp monkey model: A pet, histological and biochemical study. Neurobiol Dis 48:79–91.

Bouarab C, Thompson B, Polter AM (2019) Vta gaba neurons at the interface of stress and reward. Front Neural Circuits 13:78.

Brichta L, Greengard P (2014) Molecular determinants of selective dopaminergic vulnerability in parkinson’s disease: An update. Front Neuroanat 8:152.

Carr DB, Sesack SR (2000) Gaba-containing neurons in the rat ventral tegmental area project to the prefrontal cortex. Synapse 38:114–123.

Chaudhury D et al. (2013) Rapid regulation of depression-related behaviours by control of midbrain dopamine neurons. Nature 493:532–536.

Cho YT, Fudge JL (2010) Heterogeneous dopamine populations project to specific subregions of the primate amygdala. Neuroscience 165:1501–1518.

Chung CY, Seo H, Sonntag KC, Brooks A, Lin L, Isacson O (2005) Cell type-specific gene expression of midbrain dopaminergic neurons reveals molecules involved in their vulnerability and protection. Hum Mol Genet 14:1709–1725.

Cohen JY, Haesler S, Vong L, Lowell BB, Uchida N (2012) Neuron-type-specific signals for reward and punishment in the ventral tegmental area. Nature 482:85–88.

Dobi A, Margolis EB, Wang HL, Harvey BK, Morales M (2010) Glutamatergic and nonglutamatergic neurons of the ventral tegmental area establish local synaptic contacts with dopaminergic and nondopaminergic neurons. J Neurosci 30:218–229.

Emborg ME et al. (1998) Age-related declines in nigral neuronal function correlate with motor impairments in rhesus monkeys. Journal of Comparative Neurology 401:253–265.

Farassat N et al. (2019) In vivo functional diversity of midbrain dopamine neurons within identified axonal projections. Elife 8.

Francois C, Yelnik J, Tande D, Agid Y, Hirsch EC (1999) Dopaminergic cell group a8 in the monkey: Anatomical organization and projections to the striatum. Journal of Comparative Neurology 414:334–347.

Fu Y, Paxinos G, Watson C, Halliday GM (2016) The substantia nigra and ventral tegmental dopaminergic neurons from development to degeneration. J Chem Neuroanat 76:98–107.

Fudge JL, Kelly EA, Pal R, Bedont JL, Park L, Ho B (2017) Beyond the classic vta: Extended amygdala projections to da-striatal paths in the primate. Neuropsychopharmacology 42:1563–1576.

Galaj E et al. (2020) Dissecting the role of gaba neurons in the vta versus snr in opioid reward. J Neurosci 40:8853–8869.

Gaspar P, Heizmann CW, Kaas JH (1993) Calbindin d-28k in the dopaminergic mesocortical projection of a monkey (aotus trivirgatus). Brain Res 603:166–172.

Gaspar P, Berger B, Febvret A, Vigny A, Henry JP (1989) Catecholamine innervation of the human cerebral cortex as revealed by comparative immunohistochemistry of tyrosine hydroxylase and dopamine-beta-hydroxylase. J Comp Neurol 279:249–271.

German DC, Manaye KF (1993) Midbrain dopaminergic neurons (nuclei a8,a9, and a10): Three-dimensional reconstruction in the rat. J Comp Neurol 331:297–309.

Gorelova N, Mulholland PJ, Chandler LJ, Seamans JK (2012) The glutamatergic component of the mesocortical pathway emanating from different subregions of the ventral midbrain. Cereb Cortex 22:327–336.

Haber SN (2014) The place of dopamine in the cortico-basal ganglia circuit. Neuroscience 282C:248–257.

Haber SN, Fudge JL (1997) The primate substantia nigra and vta: Integrative circuitry and function. Crit Rev Neurobiol 11(4):323–342.

Haber SN, Fudge JL, McFarland N (2000) Striatonigrostriatal pathways in primates form an ascending spiral from the shell to the dorsolateral striatum. J Neurosci 20:2369–2382.

Haber SN, Ryoo H, Cox C, Lu W (1995) Subsets of midbrain dopaminergic neurons in monkeys are distinguished by different levels of mrna for the dopamine transporter: Comparison with the mrna for the d2 receptor, tyrosine hydroxylase and calbindin immunoreactivity. J Comp Neurol 362:400–410.

Halliday GM, Tork I (1986) Comparative anatomy of the ventromedial mesencephalic tegmentum in the rat, cat, monkey and human. J Comp Neurol 252:423–445.

Henny P, Brown MT, Northrop A, Faunes M, Ungless MA, Magill PJ, Bolam JP (2012) Structural correlates of heterogeneous in vivo activity of midbrain dopaminergic neurons. Nat Neurosci 15:613–619.

Hnasko TS, Hjelmstad GO, Fields HL, Edwards RH (2012) Ventral tegmental area glutamate neurons: Electrophysiological properties and projections. J Neurosci 32:15076–15085.

Hokfelt T, Martensson R, Bjorklund A, Kleinau S, Goldstein M (1984) Distributional maps of tyrosine-hydroxylase immunoreactive neurons in the rat brain. In: Handbook of chemical neuroanatomy, vol. Ii: Classical neurotransmitters in the cns, part i (Bjorklund A, Hokfelt T, eds), pp 277–379. Amsterdam: Elsevier.

Kim HF, Ghazizadeh A, Hikosaka O (2014) Separate groups of dopamine neurons innervate caudate head and tail encoding flexible and stable value memories. Front Neuroanat 8:120.

Kim HF, Ghazizadeh A, Hikosaka O (2015) Dopamine neurons encoding long-term memory of object value for habitual behavior. Cell 163:1165–1175.

Lammel S, Ion DI, Roeper J, Malenka RC (2011) Projection-specific modulation of dopamine neuron synapses by aversive and rewarding stimuli. Neuron 70:855–862.

Lammel S et al. (2012) Input-specific control of reward and aversion in the ventral tegmental area. Nature 491:212–217.

Lavoie B, Parent A (1991) Dopaminergic neurons expressing calbindin in normal and parkinsonian monkeys. Neuroreport 2, No. 10:601–604.

Lerner TN et al. (2015) Intact-brain analyses reveal distinct information carried by snc dopamine subcircuits. Cell 162:635–647.

Lewis DA, Campbell MJ, Foote SL, Goldstein M, Morrison JH (1987) The distribution of tyrosine hydroxylase-immunoreactive fibers in primate neocortex is widespread but regionally specific. J Neurosci 7(1):279–290.

Lowes DC et al. (2021) Ventral tegmental area gaba neurons mediate stress-induced blunted reward-seeking in mice. Nat Commun 12:3539.

Mabry SJ, McCollum LA, Farmer CB, Bloom ES, Roberts RC (2020) Evidence for altered excitatory and inhibitory tone in the post-mortem substantia nigra in schizophrenia. World J Biol Psychiatry 21:339–356.

Mailly P, Charpier S, Menetrey A, Deniau JM (2003) Three-dimensional organization of the recurrent axon collateral network of the substantia nigra pars reticulata neurons in the rat. J Neurosci 23:5247–5257.

Margolis EB, Hjelmstad GO, Fujita W, Fields HL (2014) Direct bidirectional mu-opioid control of midbrain dopamine neurons. J Neurosci 34:14707–14716.

Margolis EB, Toy B, Himmels P, Morales M, Fields HL (2012) Identification of rat ventral tegmental area gabaergic neurons. PLoS One 7:e42365.

Matsumoto M, Hikosaka O (2009) Two types of dopamine neuron distinctly convey positive and negative motivational signals. Nature 459:837–841.

Matsumoto M, Takada M (2013) Distinct representations of cognitive and motivational signals in midbrain dopamine neurons. Neuron 79:1011–1024.

McRitchie DA, Halliday GM, Cartwright H (1995) Quantitative analysis of the variability of substantia nigra pigmented cell clusters in the human. Neuroscience 68:539–551.

McRitchie DA, Hardman CD, Halliday GM (1996) Cytoarchitectural distribution of calcium binding proteins in midbrain dopaminergic regions of rats and humans. Journal of Comparative Neurology 364:121–150.

Morales M, Margolis EB (2017) Ventral tegmental area: Cellular heterogeneity, connectivity and behaviour. Nat Rev Neurosci 18:73–85.

Mugnaini E, Oertel WH (1985) An atlas of the distribution of gabaergic neurons and terminals in the rat cns as revealed by gad immunohistochemistry. In: Handbook of chemical neuroanatomy. Part i, gaba and neuropeptides in the cns (Bjorklund A, Hokfelt T, eds), pp 436–608. New York: Elsevier Science Publishers BV.

Nagaeva E et al. (2020) Heterogeneous somatostatin-expressing neuron population in mouse ventral tegmental area. Elife 9.

Nagai T, McGeer PL, McGeer EG (1983) Distribution of gaba-t-intensive neurons in the rat forebrain and midbrain. J Comp Neurol 218:220–238.

Nair-Roberts RG, Chatelain-Badie SD, Benson E, White-Cooper H, Bolam JP, Ungless MA (2008) Stereological estimates of dopaminergic, gabaergic and glutamatergic neurons in the ventral tegmental area, substantia nigra and retrorubral field in the rat. Neuroscience 152:1024–1031.

Nelson EL, Liang C-L, Sinton CM, German DC (1996) Midbrain dopaminergic neurons in the mouse: Computer-assisted mapping. J Comp Neurol 369:361–371.

Oertel WH, Tappaz ML, Berod A, Mugnaini E (1982) Two-colour immunohistochemistry for dopamine and gaba neurons in rat substantia nigra and zona incerta. Brain Res Bull 9:463–474.

Olson VG, Nestler EJ (2007) Topographical organization of gabaergic neurons within the ventral tegmental area of the rat. Synapse 61:87–95.

Olszewski J, Baxter D (2014) Cytoarchitecture of the human brainstem, 3rd, revised and extended Edition. Basel: Karger.

Omelchenko N, Sesack SR (2009) Ultrastructural analysis of local collaterals of rat ventral tegmental area neurons: Gaba phenotype and synapses onto dopamine and gaba cells. Synapse 63:895–906.

Parent A, Cote PY, Lavoie B (1995) Chemical anatomy of primate basal ganglia. Prog Neurobiol 46:131–197.

Paul EJ et al. (2018) Nnos-expressing neurons in the ventral tegmental area and substantia nigra pars compacta. eNeuro 5.

Pearson J, Goldstein M, Markey K, Brandeis L (1983) Human brainstem catecholamine neuronal anatomy as indicated by immunocytochemistry with antibodies to tyrosine hydroxylase. Neuroscience 8, No. 1:3–32.

Poulin JF, Gaertner Z, Moreno-Ramos OA, Awatramani R (2020) Classification of midbrain dopamine neurons using single-cell gene expression profiling approaches. Trends Neurosci 43:155–169.

Pristera A et al. (2019) Dopamine neuron-derived igf-1 controls dopamine neuron firing, skill learning, and exploration. Proc Natl Acad Sci U S A 116:3817–3826.

Reyes S, Fu Y, Double K, Thompson L, Kirik D, Paxinos G, Halliday GM (2012) Girk2 expression in dopamine neurons of the substantia nigra and ventral tegmental area. J Comp Neurol 520:2591–2607.

Rice MW, Roberts RC, Melendez-Ferro M, Perez-Costas E (2016) Mapping dopaminergic deficiencies in the substantia nigra/ventral tegmental area in schizophrenia. Brain Struct Funct 221:185–201.

Root DH, Estrin DJ, Morales M (2018) Aversion or salience signaling by ventral tegmental area glutamate neurons. Iscience 2:51-+.

Root DH et al. (2016) Glutamate neurons are intermixed with midbrain dopamine neurons in nonhuman primates and humans. Scientific Reports 6.

Sanchez-Gonzalez MA, Garcia-Cabezas MA, Rico B, Cavada C (2005) The primate thalamus is a key target for brain dopamine. J Neurosci 25:6076–6083.

Schein JC, Hunter DD, Roffler-Tarlov S (1998) Girk2 expression in the ventral midbrain, cerebellum, and olfactory bulb and its relationship to the murine mutation weaver. Dev Biol 204:432–450.

Selemon LD, Begovic A (2020) Reduced midbrain dopamine neuron number in the adult non-human primate brain after fetal radiation exposure. Neuroscience 442:193–201.

Seroogy K et al. (1988) A subpopulation of dopaminergic neurons in rat ventral mesencephalon contains both neurotensin and cholecystokinin. Brain Res 455:88–98.

Stark AK, Pakkenberg B (2004) Histological changes of the dopaminergic nigrostriatal system in aging. Cell Tissue Res 318:81–92.

Stauffer WR, Lak A, Yang A, Borel M, Paulsen O, Boyden ES, Schultz W (2016) Dopamine neuron-specific optogenetic stimulation in rhesus macaques. Cell 166:1564–1571 e1566.

Sulzer D, Joyce MP, Lin L, Geldwert D, Haber SN, Hattori T, Rayport S (1998) Dopamine neurons make glutamatergic synapses in vitro. J Neurosci 18(12):4588–4602.

Swanson LW (1982) The projections of the ventral tegmental area and adjacent regions: A combined fluorescent retrograde tracer and immunofluorescence study in the rat. Brain Res Bull 9:321–353.

Tan KR et al. (2012) Gaba neurons of the vta drive conditioned place aversion. Neuron 73:1173–1183.

Taylor SR, Badurek S, Dileone RJ, Nashmi R, Minichiello L, Picciotto MR (2014) Gabaergic and glutamatergic efferents of the mouse ventral tegmental area. J Comp Neurol 522:3308–3334.

Van Bockstaele EJ, Pickel VM (1995) Gaba-containing neurons in the ventral tegmental area project to the nucleus accumbens in rat brain. Brain Res 682:215–221.

van Zessen R, Phillips JL, Budygin EA, Stuber GD (2012) Activation of vta gaba neurons disrupts reward consumption. Neuron 73:1184–1194.

Wakabayashi KT et al. (2019) Chemogenetic activation of ventral tegmental area gaba neurons, but not mesoaccumbal gaba terminals, disrupts responding to reward-predictive cues. Neuropsychopharmacology 44:372–380.

Watabe-Uchida M, Zhu L, Ogawa SK, Vamanrao A, Uchida N (2012) Whole-brain mapping of direct inputs to midbrain dopamine neurons. Neuron 74:858–873.

West MJ, Gundersen HJ (1990) Unbiased stereological estimation of the number of neurons in the human hippocampus. J Comp Neurol 296:1–22.

Williams SM, Goldman-Rakic PS (1998) Widespread origin of the primate mesofrontal dopamine system. Cerebral Cortex 8:321–345.

Yamada T, McGeer PL, Baimbridge KG, McGeer EG (1990) Relative sparing in parkinson’s disease of substantia nigra dopamine neurons containing calbindin-d28k. Brain Res 526:303–307.

Yamaguchi T, Qi J, Wang HL, Zhang S, Morales M (2015) Glutamatergic and dopaminergic neurons in the mouse ventral tegmental area. Eur J Neurosci 41:760–772.

Yelnik J, Francois C, Percheron G, Heyner S (1987) Golgi study of the primate substantia nigra i. Quantitative morphology and typology of nigral neurons. J Comp Neurol 265:455–472.

Zaborszky L, Vadasz C (2001) The midbrain dopaminergic system: Anatomy and genetic variation in dopamine neuron number of inbred mouse strains. Behav Genet 31:47–59.

Zhang Y, Larcher KM, Misic B, Dagher A (2017) Anatomical and functional organization of the human substantia nigra and its connections. Elife 6.

Zhou Z et al. (2019) A vta gabaergic neural circuit mediates visually evoked innate defensive responses. Neuron 103:473–488 e476.

